# Quantitative single molecule RNA-FISH and RNase-free cell wall digestion in *Neurospora crassa*

**DOI:** 10.1101/2021.07.19.452999

**Authors:** Bradley M. Bartholomai, Amy S. Gladfelter, Jennifer J. Loros, Jay C. Dunlap

## Abstract

Single molecule RNA-FISH (smFISH) is a valuable tool for analysis of mRNA spatial patterning in fixed cells that is underutilized in filamentous fungi. A primary complication for fixed-cell imaging in filamentous fungi is the need for enzymatic cell wall permeabilization, which is compounded by considerable variability in cell wall composition between species. smFISH adds another layer of complexity due to a requirement for RNase free conditions. Here, we describe the cloning, expression, and purification of a chitinase suitable for supplementation of a commercially available RNase-free enzyme preparation for efficient permeabilization of the *Neurospora* cell wall. We further provide a method for smFISH in *Neurospora* which includes a tool for generating numerical data from images that can be used in downstream customized analysis protocols.

## 1. Introduction

*Neurospora crassa* has been in use as a model organism for the study of many important biochemical, genetic, and cell biology processes in fungi and animals for nearly 100 years (Loros, 2020; Mela et al., 2020; Riquelme et al., 2011; Selker, 2013). There are several phenomena in *Neurospora* for which the visualization and quantification of mRNA molecules could help us better understand the mechanisms fungi and other syncytial cells use to regulate protein expression, cytoplasmic organization, and mRNA trafficking. mRNA profiling has revealed heterogeneous expression of mRNA throughout mycelia of *N. crassa* (Kasuga and Glass, 2008; Mela et al., 2020). For instance, differences in gene expression can be seen even at the level of individual nuclei depending on the local cytoplasmic environment (Pieuchot et al., 2015). In addition, many biological processes are likely impacted by post-transcriptional regulation including the circadian clock which appears to require highly complex post-transcriptional regulation of mRNA. This idea is supported by the fact there are temporal delays between peak mRNA accumulation and peak protein accumulation in genes regulated by the circadian clock, in addition to gene products that are rhythmically expressed at the level of mRNA or protein but not both (Hurley et al., 2014; Hurley et al., 2018). While tools to study mRNA regulation at the cellular level have been adapted and implemented for other filamentous fungi, none have been reported in *Neurospora* (Baumann et al., 2014; Lee et al., 2016).

Fluorescence imaging of intracellular molecules in fixed *Neurospora* hyphae is challenging. While there are examples in the literature, it is not commonplace and consensus methods for chemical fixation and permeabilization have not been established (Emerson et al., 2015; Managadze et al., 2010; Riquelme et al., 2002). A major obstacle for techniques that require the use of affinity probes in intact fixed fungal cells is the presence of a cell wall, which varies in composition throughout the fungal kingdom (Bourett et al., 1998; Imdahl and Saliba, 2020; Patel and Free, 2019). Additionally, we have found that mature, fixed *Neurospora* hyphae are recalcitrant toward adherence to glass or other surfaces commonly used for light microscopy. While there are several lytic enzyme preparations commercially available for permeabilization of fungal cell walls, they are not universally effective due to compositional differences in walls amongst fungi. Additionally, the enzymes tend to be harvested from cellular extracts that are enriched for fungal lytic enzyme activity by size or charge based chromatographic methods, but are not necessarily pure or free from contaminating nuclease activity, which is particularly important for the study of RNA.

Here we describe a method for liquid culture growth, chemical fixation, and digestion of the *N. crassa* cell wall in RNase-free conditions. RNase-free cell wall digestion employs a recombinant chitinase, for which we also provide a method for expression and purification. Finally, we have adapted single molecule RNA-FISH (smFISH) for use in *Neurospora* from protocols developed for other fungi (Lee et al., 2016; Raj et al., 2008). In total, this approach enables the quantitative analysis of mRNA localization and abundance in *Neurospora* opening up the ability to ask fundamental questions about RNA regulation in this key model system.

## 2. Recombinant *Bacillus licheniformis* chitinase as a tool for RNase-free cell wall digestion of *N. crassa*

A critical first step in FISH protocols for fungi is permeabilizing the cell membrane and walls sufficiently to allow large, nucleic acid probes to enter the cell. The *Neurospora* cell wall is comprised of β-Glucans, chitin, galactomannan, and proteins. A compact layer of chitin is situated directly against the plasma membrane (Free, 2013; Verdín et al., 2019). Zymolyase is a common enzymatic preparation used for digestion of cell walls in fungi (Lee et al., 2016; Li and Neuert, 2019). While highly purified Zymolyase-100T is RNase-free (amsbio, cat. # 120493-1), it has not been shown to be effective in degrading the *Neurospora* cell wall and it does not have chitinase activity. In preliminary studies, we found that Zymolyase did have some effect on the *Neurospora* cell wall, but it was not effective in permeabilizing them (data not shown). To overcome this challenge, we cloned, expressed, and purified a *Bacillus licheniformis* chitinase. Chitinase from *B. licheniformis* has been reported to affect cell wall composition in several filamentous fungi, while being easily expressed and purified from BL21 *Escherichia coli* (Gomaa, 2012; Songsiriritthigul et al., 2010; Yamabhai et al., 2008). Here we report that this recombinant chitinase can be purified as an active, RNase-free, enzyme that significantly improves cell wall digestion when used with Zymolyase-100T.

### 2.1. Cloning, expression, and purification of *B. licheniformis* chitinase

We acquired a live *B. licheniformis* culture from Ward’s Science (Item # 470179-060), from which a single colony was isolated and expanded for genomic DNA extraction using Omega Bio-Tek E.Z.N.A. Bacterial DNA Kit (Item # D3350-00). A DNA fragment was amplified that corresponds to the ChiA expressing gene described by Yamabhai and colleagues (NCBI accession # AAU21943) using Phusion Flash (Thermo Scientific Item # F548L). Primers were designed to insert the gene of interest into pET16better (provided by the lab of Dean Madden at Dartmouth) using NEBuilder DNA assembly (New England BioLabs Item # E2621S). The resulting plasmid contains a T7 promoter that drives expression of ChiA containing a 10XHis tag followed by a HSV3c cleavage site at the N terminus (Figure 1A).

**Figure 1.**
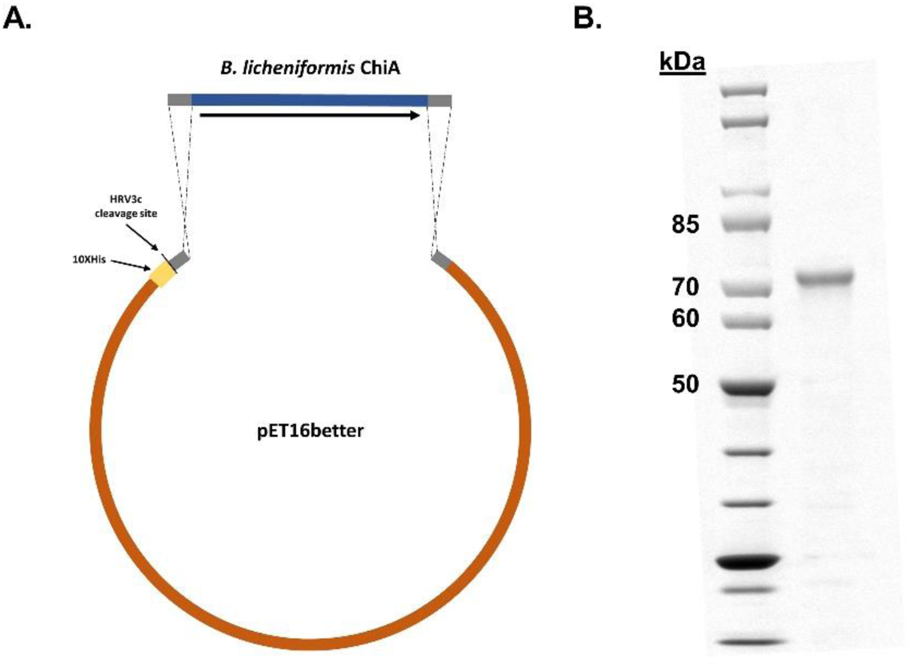
*Bacillus licheniformis* Chitinase (ChiA) was expressed in *E. coli* and purified. A) *B. lichenifomris* ChiA was amplified from genomic DNA by PCR and assembled into pET16better by Gibson assembly. B) Chitinase (68.9 kDa) was expressed in *E. coli* and purified to ≥ 90% on an FPLC by immobilized metal affinity chromatography and size exclusion chromatography.

We selected SHuffle^®^ T7 Express Competent *E. coli* (NEB Item # C3029J) as the expression strain because the ChiA protein contains 6 cysteine residues. This strain expresses the chaperone DsbC, which should aid in proper disulfide bond formation. Plasmid DNA was transformed into competent *E. coli.* A single colony from overnight LB-agar plates containing 100 μg/mL carbenicillin was used to grow a 50 mL overnight starter culture in Terrific Broth containing 100 μg/mL carbenicillin (TB + crb). Four liters of TB + crb separated into four 2-liter baffled flasks were warmed to 37 °C. Pre-warmed liters of TB + crb were inoculated with 10 mL of starter culture that had been shaking at 250 RPM and 37 °C for 16 hours. The cultures were shaken at 250 RPM and 37 °C and OD600 measurements were taken regularly until they reached a value of ~0.4, at which point they were moved to shaking at 250 RPM and 16 °C until they reached an OD600 between 0.6 and 0.8. Recombinant chitinase expression was induced with 1 mM IPTG and cultures were left to shake at 250 RPM and 16 °C for 18 hours.

Bacterial cultures were pelleted using a tabletop centrifuge equipped with a swinging arm rotor at 3000 g for 10 minutes and the supernatant was discarded. Pellets were resuspended in purification Buffer A (50 mM HEPES pH 7.4, 1M NaCl, 20 mM Imidazole, 10% glycerol, and 1 mM TCEP) with 2X Roche cOmplete™, EDTA free protease inhibitor cocktail (MilliporeSigma Item # 11873580001) using 1 ml Buffer per 0.25 g wet pellet mass. Cells were lysed by 4 passes through a LM10 Microfluidizer™ Processor. Lysates were cleared using a Beckman Optima L-70 preparative ultracentrifuge, with a 45 Ti rotor at 42000 RPM for 1 hr at 4 °C. Supernatant was filtered through a 0.45μm syringe filter. Immobilized metal affinity chromatography (IMAC) was carried out on a Cytiva Äkta FPLC at 4 °C using a pre-packed 5 mL HisTrap HP column (Cytiva Item # 17524802). NiNTA purification was carried out with two buffers, Buffer A and Buffer B (50 mM HEPES pH 7.4, 1M NaCl, 500 mM Imidazole, 10% glycerol, and 1 mM TCEP) using the following protocol. The HisTrap HP column was primed and equilibrated with 5 column volumes (CV) of Buffer A, followed by 5 CV Buffer B, then 10 CV Buffer A at a flow rate of 5 ml/min. Cleared lysate was added to the column using a sample pump at a rate of 2.5 ml/min. The column was washed with 5 CV Buffer A at a rate of 5 ml/min. Elution was carried out step wise in 5 CV increments, using 3 increasing concentrations of Buffer B (10%, 20%, and 100%) at a rate of 5 ml/min. 5 ml fractions were collected and retained for further analysis. The chromatogram shows a strong A280 peak in the fractions corresponding to 100% Buffer B (12-16) (Supplemental Figure 1). Fractions 12-16 were pooled and Triton^®^X-100 was added to a final concentration of 0.5% to facilitate removal of interacting proteins. A second purification step using size exclusion chromatography (SEC) was implemented to remove excess Imidazole, Triton^®^X-100, and isolate the recombinant chitinase. SEC was also carried out on a Äkta FPLC at room temperature using a HiLoad 26/600 Superdex 200 pg column (Cytiva Item #28989335). Equilibration of the column with RNase-free SEC Buffer (50 mM HEPES pH 7.4, 150 mM NaCl, 10% glycerol, and 1 mM TCEP) was carried out by flowing buffer through the column at a rate of 2.6 ml/min. Protein sample was added to the column using a capillary loop at a rate of 2.6 ml/min. Elution was carried out with 1.3 CV SEC Buffer. 5 ml fractions were collected after 0.2 CV in 13 × 100 mm glass culture tubes that had been baked in a laboratory oven at 200 °C for 4 hours to avoid RNase contamination. Protein was quantified using Pierce^®^ 660 nm Protein Assay (Thermo Scientific Item # 1861426) and absorbance at 280 nm with the expected molecular weight of 68.93 kDa and predicted extinction coefficient of 154,240 M^-1^ cm^-1^. Protein parameter predictions were generated with the Expasy ProtParam tool (Gasteiger et al., 2005). 1 μg purified chitinase was run on a 4-12% Invitrogen Bolt Bis-Tris Plus gel (Invitrogen Item # NW04125BOX) and stained with Blazin′ Blue™ Protein Gel Stain (GoldBio Item # P-810-1). Visualization of the gel shows a single prominent band at the expected size (Figure 1B).

### 2.2. Activity analysis of recombinant chitinase

Chitinase activity was assessed by performing a continuous colorimetric assay adapted from a previously described assay (Lobo et al., 2013). 20 μg/ml recombinant chitinase was incubated with 1 mg/ml 4-Nitrophenyl β-D-N,N′,N″-triacetylchitotriose (MilliporeSigma Item # N8638) for 30 minutes at 37 °C on a Synergy Neo2 multi-mode plate reader. Release of 4-nitrophenol from the substrate was measured by absorbance at 410 nm every 2.5 minutes. First degree polynomial regression was performed on A410 values of a standard curve of known 4-nitrophenol values measured on the same plate as the chitinase assay (Supplementary Figure 2). A410 values of 4-nitrophenol released in the chitinase assay were converted to concentration using the standard curve and plotted (Figure 2A). Polynomial regression of the chitinase activity assay results provided a reaction rate of 0.009922 μmol/ml/min. The reaction rate was converted to enzyme units defined as the amount of enzyme required to catalyze 1 μmol of substrate per minute (1 U = 1 μmol/min). We have determined the specific activity of this chitinase to be 2.02 U/mg at 37 °C. Reported activity values for *B. licheniformis* chitinase A vary widely. This is thought to be because of the water insoluble nature of the natural target, chitin, and difficulty determining its precise molecular weight. Various synthetic substrates such as 4-nitrophenyl conjugated chitotrise and chitobiose have been used, as well as partially soluble colloidal chitin (Menghiu et al., 2019). Much of the research surrounding *B. lichenifomis* chitinase A involves thermophilic environmental isolates, in which high temperatures were used to determine activity that are not relevant to the experiments we intend to conduct (Asmani et al., 2020; Laribi-Habchi et al., 2015). At 37 °C, specific activity of this chitinase has been reported as 0.294 U/mg with *p*-NP-chitobiose as the substrate. The same group also performed kinetic analysis at 37 °C using colloidal chitin as the substrate and standard curves of *N*-acetylglucosamine or di-*N*-acetyl chitobiose and found the specific activity to be 1.5 U/mg and 5.11 U/mg respectively (Songsiriritthigul et al., 2010).

**Figure 2.**
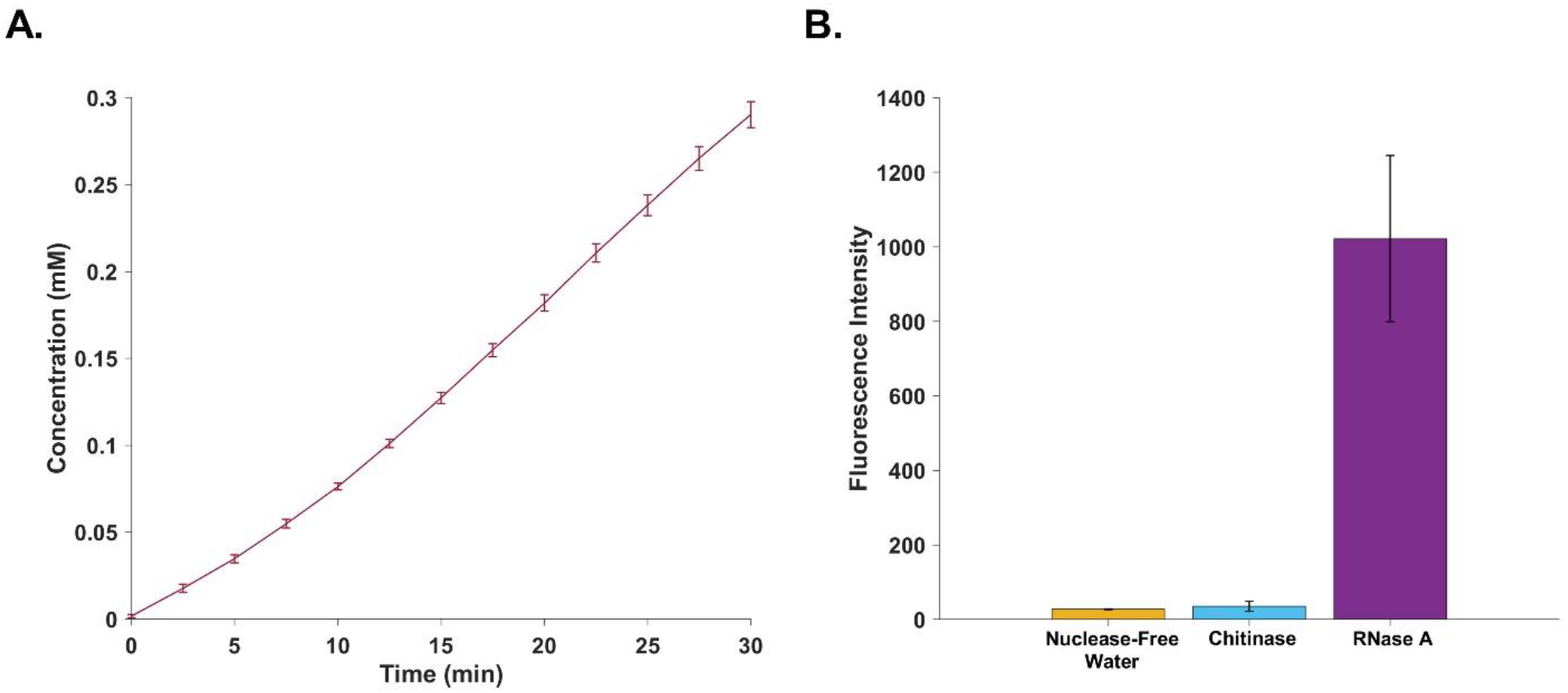
Purified chitinase is active and free of RNase. A) Purified chitinase (290 nM) was incubated at 37 °C with 4-Nitrophenyl β-D-N,N′,N″-triacetylchitotriose (1.34 mM). Absorbace at 410 nm was measured every 2.5 minutes. Concentration of 4-nitrophenol released from the substrate was determined by a standard curve of known values. B) RNase activity was assessed using the RNase Alert^®^ Lab Test Kit. Nuclease-Free Water, chitinase (0.04 U/ml), and RNase A (0.03 U/ml) were incubated at 37 °C for 30 min and measured with a fluorometer.

Assessment of RNase activity in the purified chitinase at a concentration we have found is sufficient for smFISH, 0.02 mg/ml, was carried out using a RNase Alert^®^ Lab Test Kit v2 (Invitrogen Item # 4479768). Nuclease-Free Water and 0.03 U/ml RNase A provided in the kit were used as negative and positive controls, respectively. Reactions were carried out in triplicate for all three samples following the manufacturer’s instructions. Test samples were incubated at 37 °C for 30 min with the substrate, which is a small RNA molecule with a fluorophore conjugated to one end and quencher conjugated to the other. Upon RNA cleavage the fluorophore is released, which can be measured and correlated to RNase activity. According to the manufacturer, as little as 0.3 pg of RNase A can be detected in each 50 μl reaction. The mean fluorescence intensity values for water, chitinase, and RNase A were 27, 35.3, and 1022 respectively. The values of chitinase and the negative control were not statistically different. According to the kit manufacturer, samples with fluorescence intensity values 2- to 3-fold greater than the negative control are considered contaminated with RNase. However, samples that are contaminated with RNase typically yield fluorescence intensity values 20- to 100- fold higher than the negative control, as seen in the positive control. Results of this assay verify that our purified recombinant chitinase is free of RNase contamination and is therefore suitable for use in smFISH experiments (Figure 2B).

## 3. Single molecule RNA-FISH in *Neurospora crassa*

smFISH is a powerful tool that has been validated and used in many systems to visualize mRNA in cells but underutilized in filamentous fungi. A major reason for this is likely the difficulty presented by cell wall permeabilization and maintaining cellular integrity throughout fixed-cell imaging procedures. Here, we present a protocol which was adapted from previous work in the filamentous fungus *Ashbya gossypii* (Dundon et al., 2016; Lee et al., 2016; Lee et al., 2013). Taken together, smFISH protocols for *Ashbya* and *Neurospora* are likely to help facilitate easier adaptation to other filamentous fungal systems.

### 3.1. Culturing of hyphae and fixation

Cultures are grown overnight in liquid media starting with conidia. To generate conidia, inoculate agar slants made with Vogel’s minimal medium (Vogel, 1956) with conidia from frozen slants or glycerol stocks. After 5-7 days of growth at 25 °C, suspend conidia in sterile water and briefly pellet in a microcentrifuge. Discard the supernatant containing hyphal debris. Resuspend conidia in water and filter through cheesecloth. Inoculate 27 ml of liquid Vogel’s minimal medium pH 6.5, containing 2% glucose and 0.2% polyacrylic acid with no more than 1 x 10^6^ conidia/ml. Shake liquid cultures as previously described (Kelliher et al., 2020). The primary benefit of Kelliher’s method for imaging studies on the temporal scale we are interested in is the use of 0.2% polyacrylic acid in the medium to prevent hyphal fusion and maintain a dispersed culture. Use of liquid medium without polyacrylic acid gives rise to large aggregates of fused hyphal tissue that cannot be used for downstream imaging protocols, particularly when hyphae are incubated for multiple days for studies of the circadian clock, the primary focus of our lab.

For circadian biology experiments, conidial suspensions are incubated at 25 °C, shaking at 100 RPM in constant light for 6 hours and moved to constant darkness at the same temperature with continued shaking until the desired circadian timepoint is reached. Kelliher et al., 2020 found that glucose is fully depleted from the cultures after about 48 hours, so the total age of our shaking cultures is typically between 22 and 42 hours. When desired culture growth has been achieved, fix cultures by adding 3 ml of 37% formaldehyde, for a final concentration of 3.7% formaldehyde (10% formalin). Continue shaking the cultures containing formalin for 45 – 60 min. Fixation times less than this lead to poor structural preservation. To avoid RNase contamination, all work after fixation should be carried out under sterile, RNase-free conditions. Transfer 5 ml of fixed culture to an RNase-free 5 ml conical tube (VWR Item # 10002-731) using an RNase-free serological pipette, and centrifuge at 1000g for 2 min at 4 °C in a tabletop centrifuge with a swinging bucket rotor. Aspirate supernatant, making sure not to disturb the loose pellet. Resuspend the pellet in ice-cold DEPC treated buffered sorbitol (100 mM Potassium Phosphate pH 7.4 and 1.2 M Sorbitol). Wash resuspended hyphae for 5 min on an end over end rotator at 4 °C. Repeat the previous steps beginning with centrifugation twice for a total of 3 washes. At this point, fixed hyphae can be stored in buffered sorbitol for a few days at 4 °C or carried through immediately to cell wall digestion.

### 3.2. Cell wall digestion and permeabilization

Cell wall digestion solution (100 mM Potassium Phosphate, pH 7.4, 1.2 M Sorbitol, 250 U/ml Zymolyase 100T, 20 μg/ml recombinant chitinase, and 5 mM Ribonucleoside Vanadyl Complex (New England Biolabs Item # S1402S)) should be prepared fresh for each experiment. Add 0.5 ml of fixed hyphae to 2.5 ml digestion solution in a new RNase-free 5 ml conical tube and incubate at 37 °C on a nutating mixer. Cell wall digestion can be assessed over time using calcofluor white (CFW). CFW is a fluorescent dye widely used to image the cell wall of various microbes and plants ( (Darken, 1961; Harrington and Hageage Jr, 2003; Hoch et al., 2005; Ursache et al., 2018). Observation of the cell wall with CFW is facilitated by its binding to 1-3 β and 1-4 β polysaccharides.

Inconsistent cell wall digestion in early attempts to adapt this protocol for *Neurospora* led us to develop the recombinant chitinase described above. Here, we compare undigested hyphae to hyphae digested with chitinase and Zymolyase, as well as Zymolyase alone, expecting to see fluorescence intensity from CFW staining diminish over time as the cell wall is degraded by the lytic enzymes. This method is also used in the smFISH protocol to assess cell wall degradation over time. Cell wall composition and therefore the time required for digestion can vary between different strains and different stages of the circadian cycle. Therefore it is important to monitor this process each time the procedure is carried out. Remove 100 μl of fixed culture from the digestion solution, pipette into a 1.5 ml microcentrifuge tube and mildly vortex with 1 μl 5 mM CFW (Biotium Item # 29067). Pipette 20 μl of CFW stained culture to a plain glass microscope slide, overlay the stained hyphae with a 24 x 50 mm coverslip, and tack the corners down with clear nail polish. We carry out imaging for this process with a Nikon Ti-E widefield fluorescence microscope equipped with a Lumencor Sola Light Engine and an Andor Zyla 4.2 camera, using Chroma filter set 49028 (EX 395/25x and EM 460/50m) and a 60X oil immersion objective. However, any microscope capable of fluorescence excitation at 350 nm and emission detection at 450 nm should be sufficient. The cell wall of *Neurospora* can be clearly observed as a bright fluorescent outline of the hyphae in undigested cells. By 60 min of incubation with Zymolyase and Chitinase, nearly all of the signal in wild-type samples (strain *74-OR23-1VA* (FGSC # 2489) grown for 24 hours) is lost as observed by the human eye. Hyphae incubated with Zymolyase alone, however, only show nominal qualitative signs of degradation (Figure 3A).

**Figure 3.**
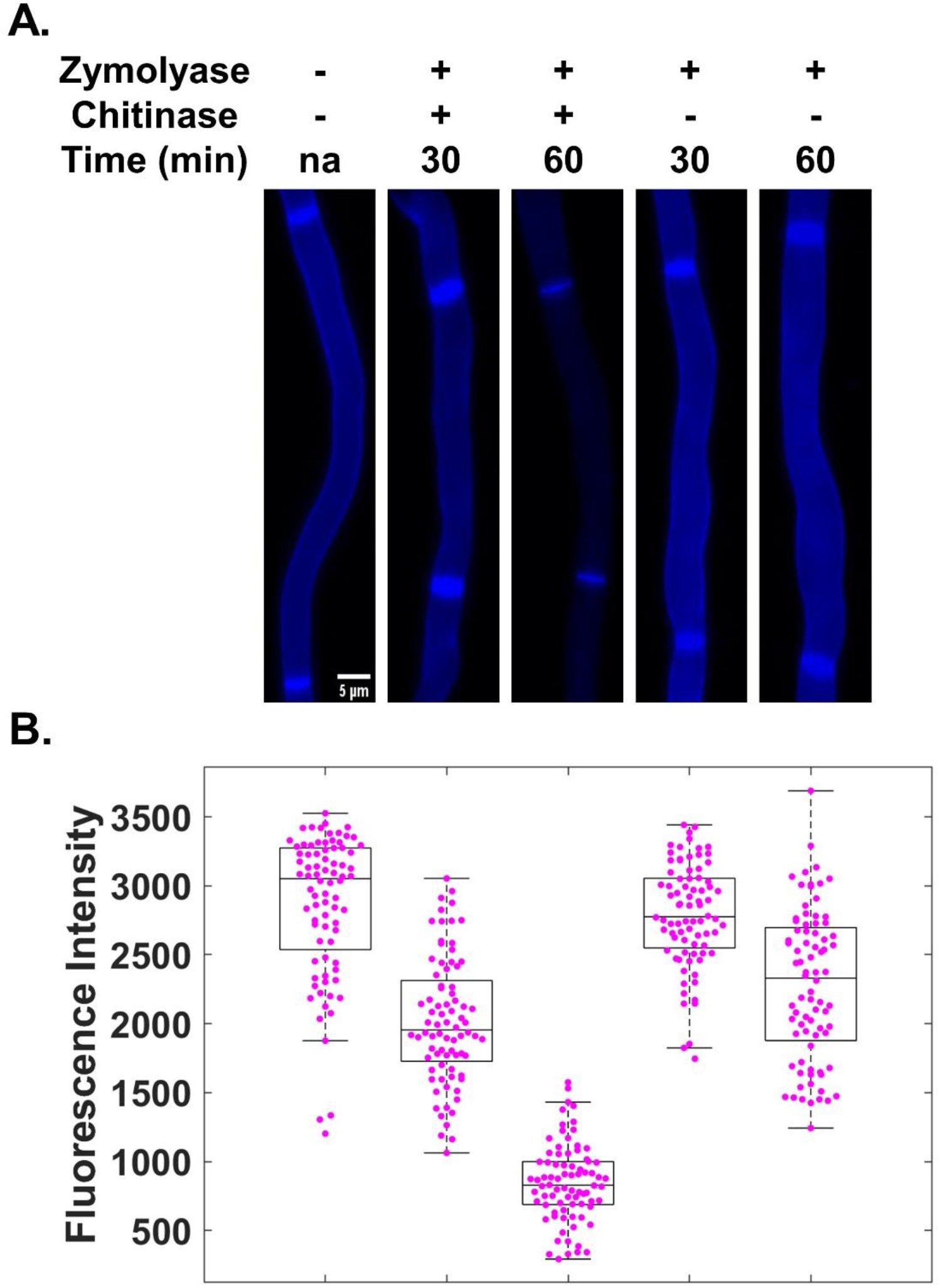
Chitinase provides more efficient, consistent, and complete cell wall digestion than Zymolyase 100T alone. A) Representative fixed *N. crassa* hyphae stained with Calcofluor White (50 μM) after incubation at 37 °C for the indicated amount of time with Zymolyase 100T or Zymolyase 100T plus Chitinase. B) Quantification of fluorescence intensity normalized to background fluorescence, indicating concentration of chitin and other fibrillar polysaccharides. 80 measurements were taken from 20 images for each condition.

Quantification of imaging data for the experiment in figure 3 was carried out by conducting line scans perpendicular to the hypha, using a uniform line width that extended beyond the borders of the hyphae at 4 random positions (avoiding the highly stained septa) for each of 20 hypha from each condition. Assuming the maximum intensity of a given line is at the cell wall where 1-3 β and 1-4 β polysaccharides are concentrated, the maximum intensity values were normalized by subtracting the minimum value, which is assumed to be background from the region of the line outside the cell. Boxplot representation of the data with overlayed data points shows considerably improved performance of Zymolyase when combined with chitinase over Zymolyase alone (Figure 3B). Statistical analysis of these data by one-way ANOVA confirmed significant variation between the different conditions (P < 3.43 × 10^-110^). Multiple comparison analysis by Tuckey’s HSD revealed that the mean value of 30 minutes of incubation with Zymolyase alone is not statistically different from the undigested sample, while all other pairwise comparisons have P<.005.

After cell wall digestion is complete, with approximately 80-90% of observed cells showing no strong fluorescence from CFW staining at the cell wall, wash hyphae 3 times in the same manner as previously described after fixation. After the final wash, centrifuge hyphae once more and aspirate the supernatant, leaving about 0.2 ml of buffered sorbitol in the tube, which can be approximated by the markings on the tube. Membrane permeabilization, lipid extraction, and protein denaturation is carried out by adding ice-cold 70% EtOH to a final volume of about 3.0 ml. This step must be carried out slowly and step wise. Addition of ethanol too quickly tends to cause aggregation of the hyphae. In samples that have aggregated, we have found that attempting to carry out the rest of the procedure results in poor hybridization, bad signal to noise, and uneven mounting due to large aggregates of hyphae. Aggregation causes hyphae to exist in such a broad range of focal planes, that segmentation and quantification is virtually impossible. If aggregation occurs at this step, we highly recommend discarding the cells and returning to the cell wall digestion step with a new aliquot of fixed hyphae. Hyphae should be kept in EtOH at least overnight at 4 °C before proceeding and can be stored in this manner for at least a week.

### 3.3. Hybridization of fluorescent mRNA oligonucleotide probes

After permeabilization is complete, pipette 0.5 ml of hyphae stored in 70% EtOH into a new RNase-free 5 ml conical tube, add 4.5 ml wash buffer (2X RNase-free SSC and 10% deionized formamide), mix for 5 min on a nutating mixer at room temperature, and centrifuge for 2 minutes at 1000g. This is repeated twice for a total of 3 washes. After the final wash and centrifugation, aspirate supernatant to about 0.2 ml and resuspend hyphae by gentle flicking. It is important to start with an excess of hyphae in the tube at the beginning of these washes. At this step, hyphae tend to adhere to the walls of the plastic tube and considerable loss of sample is observed. Gentle vortexing to remove hyphae from the walls of the tube does not seem to have deleterious effects, but too much aggressive handling can lead to severe fragmentation of hyphae. We have also found that use of 1.5 ml low adhesion tubes (SSIbio item # 1210-00) with appropriate volume adjustments can dramatically reduce loss of hyphae when used with a centrifuge equipped with a swinging bucket rotor. Fragile fixed and permeabilized hyphae do not tend to pellet well in a fixed angle centrifuge at speeds low enough to not damage the cells.

Fresh hybridization medium should be prepared for each experiment by adding deionized formamide to Stellaris Hybridization Buffer (Biosearch Technologies Item # SMF-HB1-10) for a final concentration of 10%. We purchase Stellaris smFISH probes from Biosearch Technologies and use their RNA FISH Probe designer to optimize the position and number of probes. This program uses the DNA sequence of an mRNA of interest to optimize the positioning of 25 – 48 20-mer oligonucleotides with a conjugated fluorophore across the full length of the molecule. We have observed good results with Quasar^®^ 670 and CALFluor 590 conjugated probes, but only limited success with Quasar^®^ 570 conjugated probes. Dyes with excitation spectra closer to green wavelengths seem to have more trouble with signal to noise, presumably due to cell autofluorescence. Optimization needs to be carried out for each probe set, but we have found that a final dilution of 1:100 of the stock preparation Stellaris recommends is a good place to start.

Incubate fixed and permeabilized hyphae that have been gently mixed with smFISH probes and hybridization buffer at 37 °C for 16 – 18 hours.

### 3.4. Nuclear staining and mounting of hyphae

Upon completion of hybridization, add 4 ml of wash buffer pre-warmed to 37 °C to each hybridization reaction and mix by inversion. Return samples to 37 °C and incubate for 30 minutes. After this initial wash step, centrifuge hyphae at 1000 g in a tabletop centrifuge equipped with a swinging bucket rotor for 2 minutes and aspirate supernatant, leaving about 0.2 ml. Resuspend hyphae by gentle flicking and pipette into silicone wells (Electron Microscopy Sciences Item # 70325-52) on Superfrost^®^Plus slides (VWR Item # 48311-703). The slides used for this are positively charged during the manufacturing process. This facilitates adherence of hyphae to the glass surface, to preserve samples in subsequent wash steps. Place slides at 37 °C for about 20 – 30 minutes while hyphae settle. In some cases, the hyphae will not settle, and extra care is required in the following steps. Typically, when hyphae contact the glass surface, they will stay in place for the remainder of the procedure. Carefully aspirate buffer from the well and replace with wash buffer containing 0.02 mM Hoechst 33342. Incubate samples in this solution at room temperature, protected from light, for 15 minutes followed by careful aspiration of the buffer. A final wash step is performed by pipetting 250 μl wash buffer into the well and incubating at room temperature for 5 – 10 minutes.

After washing and nuclear staining is complete, carefully aspirate buffer from the well, gently remove the silicone well from the slide, and aspirate any residual liquid. Pipette 10 μl of ProLong™ Glass Antifade Mountant (Invitrogen Item # P36984) to the center of the sample and overlay with a sterile #1.5 22 x 22 mm glass coverslip. Selection of cover glass should consider the calibration of the objective to be used for imaging and may be different for different systems. Mounted slides are left face up in the dark for several hours to allow any air bubbles to resolve. The slides can be covered by a Kimwipe and a flat surfaced object such as a hard bound book to assist in removal of bubbles. To allow the mounting medium to fully harden, slides are placed in a rack within an airtight container with desiccant overnight at 4 °C. Slides should be imaged as soon as possible, but can be stored at 4 °C, protected from light for at least a few days without any loss of signal

### 3.5. Imaging and quantification of mRNA and nuclei

We have successfully imaged smFISH samples using widefield and spinning disk confocal systems. Widefield images typically require quite a bit more post-processing because of out of focus light, so we typically use a spinning disk confocal system. Our preferred system is a Nikon Ti-E based inverted microscope equipped with a Yokogawa CSU-W1 confocal scanner unit and dual Photometrics Prime BSI sCMOS cameras. We use a 60X oil immersion lambda series objective with a numerical aperture of 1.4. To gather 3D information, we acquire z-stacks with oversampling at a rate of 0.7 over the Nyquist rate to improve resolution of discrete mRNA puncta. Using this technique to visualize the G1/S cyclin *cln-1,* we can clearly visualize discrete puncta throughout the hyphae in the smFISH channel, which are qualitatively consistent with previous studies of this cyclin in *Ashbya* (Lee et al., 2013) (Figure 4a). Nuclear staining is consistent with fixed-cell imaging conducted by our group and others (Emerson et al., 2015; Riquelme et al., 2002) (Figure 4b). A composite image shows bright foci, apparently co-localized with nuclei, which is consistent with previous reports of nuclear transcriptional hotspots, indicating adequate permeabilization of intracellular structures (Lee et al., 2016)(Figure 4c).

**Figure 4.**
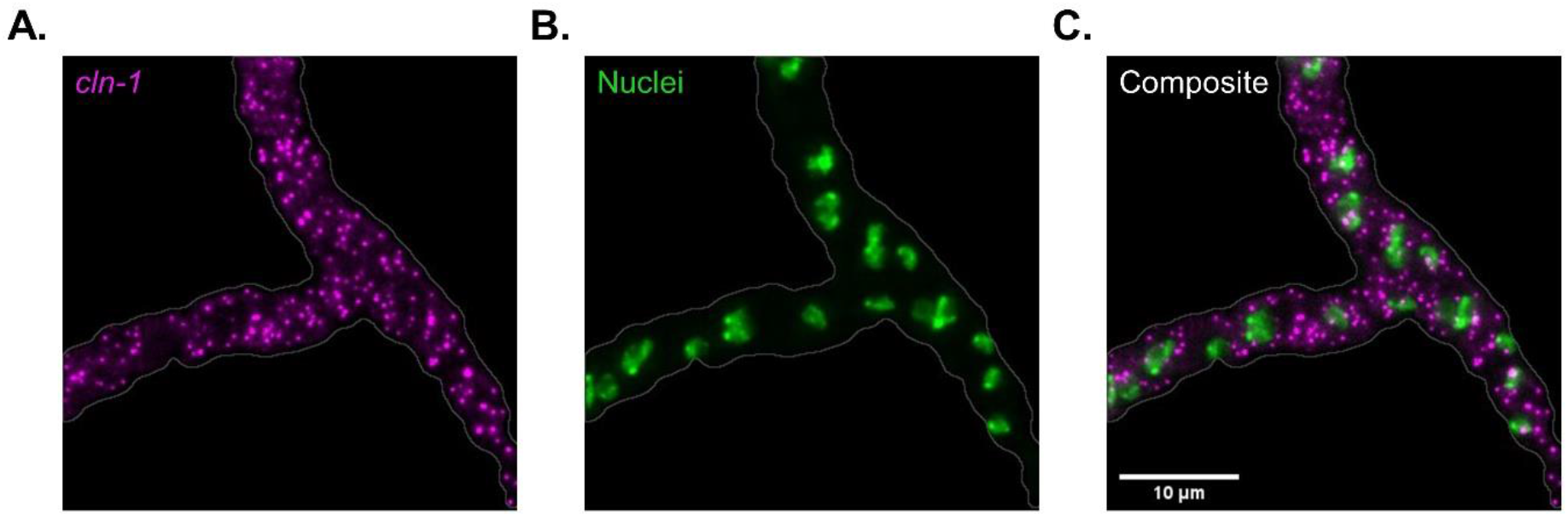
smFISH can be used to visualize mRNA in *Neurospora crassa*. This maximum intensity z-projection shows *cln-1* mRNA, which has been hybridized to 48 20-mer oligonucleotides conjugated with Quasar^®^670 in fixed wild-type *Neurospora crassa 74-OR23-1VA* (FGSC # 2489) hyphae (A). Nuclei are visualized using Hoechst 33342 stain (B). The channels are false colored magenta and green to assist in differentiation between the channels in the composite image (C). A gray outline of the hyphae is provided to delineate the boundaries of the cell.

Quantification of images can be carried out with open-source or commercially available software. Here, we provide a method and macro for 3D image detection using the open source software FIJI adapted from a previously published method (Lee et al., 2016). This macro processes all 2 channel images in a folder with real-time user interaction. First, it separates the channels (Figure 5 A). Next, it asks the user to define a segmenting threshold using the 3D Objects Counter plugin for each channel separately. This plugin produces a map of detected objects and centroids which can be inspected by the user to ensure correct segmentation in real time (Figure 5 B and C) (Bolte and Cordelières, 2006). Statistics of fluorescence intensity and object 3D location are exported to a .csv file and automatically saved in a designated folder. Finally, a Z-projection is used to create a mask at the widest point of the hyphae and exports the x-y coordinates of the mask to a .csv file. This 2 dimensional mask can be converted in downstream applications to 3D assuming a cylindrical shape as described previously (Lee et al., 2016).

**Figure 5.**
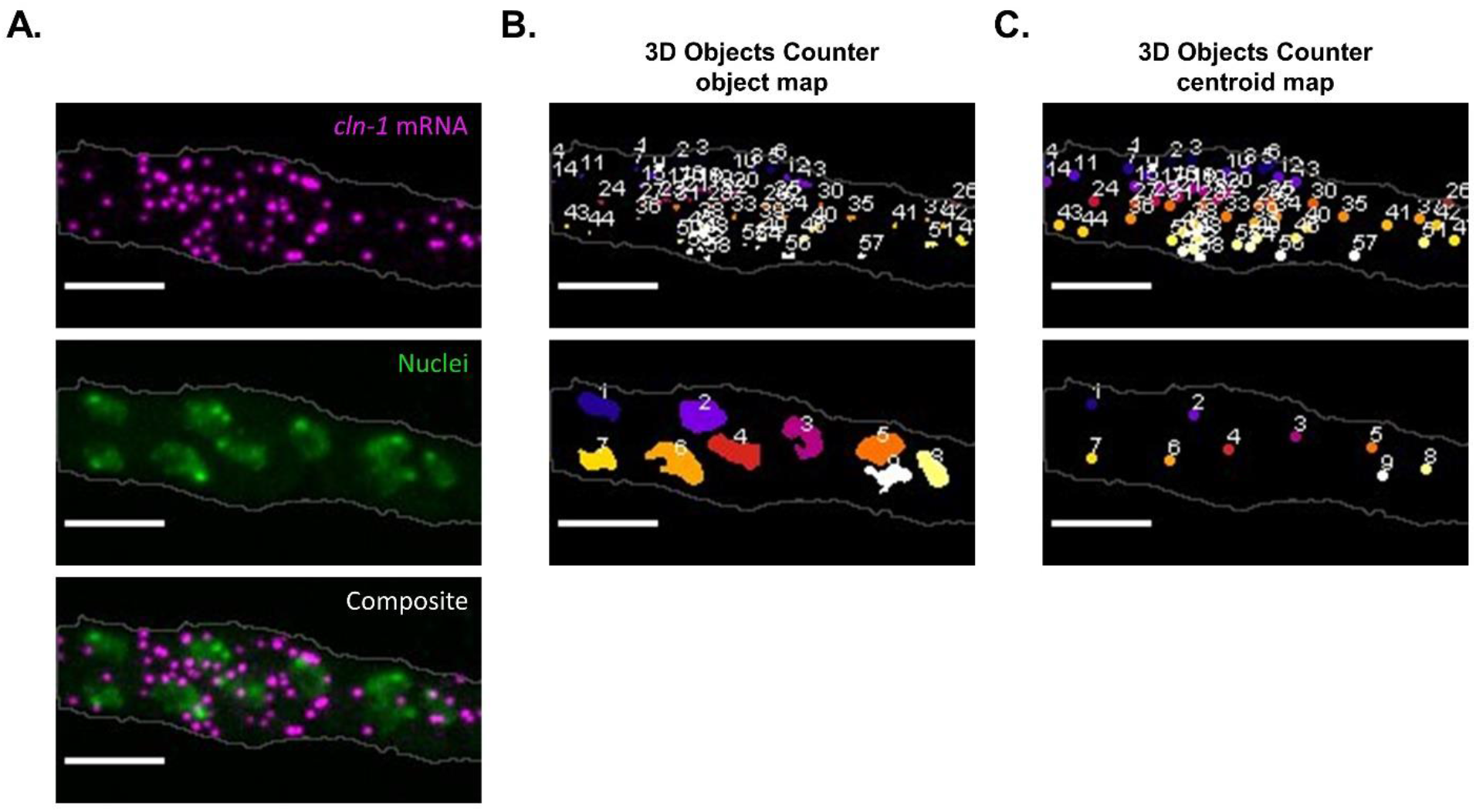
mRNA can be quantified in *N. crassa* using image analysis software. A) Maximum intensity z-projection of fixed and permeabilized *N. crassa* hyphal segment hybridized with custom *cln-1* specific probes conjugated with Quasar670 and nuclei stained by Hoechst 33342. B) Map of objects detected by the 3D Objects Counter FIJI plugin. C) Map of centroids determined by the 3D Objects Counter FIJI plugin.

## 4. Conclusions

Difficulties surrounding reliable fixation and permeabilization for fluorescence imaging methods has limited the application of these techniques in filamentous fungal research compared to more tractable systems like yeast, insects, and mammals. With recent advancements in single molecule imaging and super resolution microscopy techniques that require fixed cells, improved methods are necessary to enable biological discovery in filamentous fungi. The introduction of the recombinant chitinase demonstrates that despite diversity in cell wall composition, customization of techniques to specific fungi is possible and well-defined enzyme mixtures can be created to alleviate concerns surrounding potential contaminating factors such as RNase or other undesirable enzymes. *Neurospora* in particular has a wealth of transcriptomic data and biochemical characterization of several RNA binding proteins for which the techniques described here can be useful for probing at the cellular level (Caster et al., 2016; Craig et al., 2015; Herold and Yarden, 2017; Hurley et al., 2014; Liu et al., 2017; Wu et al., 2014).

Several differences between *Neurospora* and *Ashbya* led to the need for adaptations to the protocol for sample preparation and analysis. *Neurospora* grows more quickly than *Ashbya,* branches less, has longer hyphae that are more variable in diameter, more nuclei that are not evenly distributed in the cytoplasm, and it creates an interconnected network via hyphal anastomosis. Liquid growth conditions were optimized to minimize anastomosis and ensure a suspension culture that can be pipetted after fixation. Cell wall digestion was customized to efficiently permeabilize the cell wall. Addition of ethanol was changed to slow, step wise addition because of problems with aggregation that were encountered. All centrifugations were carried out in a centrifuge with swinging bucket rotors because *Neurospora* would not pellet in fixed angle centrifuges at speeds that did not cause sheering of delicate fixed/permeabilized hyphae. Final washes were carried out on a charged slide to facilitate hyphae settling on the surface and adhering to improve image acquisition. The mounting method for *Ashbya* allowed long *Neurospora* hyphae to undulate throughout the mounting medium, and individual hyphae were rarely in the same focal plane, which adversely affected segmentation.

As with any technique, there are some limitations of smFISH for fungal systems. Heterogeneity throughout mycelial systems is a well-documented and important phenomenon (de Bekker et al., 2011; Mela et al., 2020; Wösten et al., 2013; Zacchetti et al., 2018). In fungi like *Neurospora* that are highly buoyant and do not adhere well to surfaces commonly used for microscopy, investigation of fused mycelial systems would be extremely challenging. We have observed heterogeneity in signal concentrations throughout cultures (Supplementary Figure 3). However, a hallmark of the polyacrylic acid containing growth medium we use is that hyphae stay dispersed and tend not to fuse. Liquid medium growth is commonplace for the study of circadian rhythms and other phenomena in *Neurospora.* Therefore, this technique should provide results that are consistent with previous studies. While this is beneficial for several reasons and appears to give us genuine biological insights, the cell biology of naturally growing fungi on a solid substrate might be quite different. Another limitation is that fixed hyphae tend to undergo plasmolysis. While this can be avoided by imaging of cells that look the most intact, it is possible that morphological changes due to fixation might not be fully representative of the architecture of living cells.

There are many potential uses for this technique in *Neurospora* and other filamentous fungi to understand the cellular context of RNA regulation as it relates to trafficking, localization, spatial patterning, and relationships between RNA and RNA binding proteins. Here, we have developed a method that allows for effective chemical fixation and reliable cell wall removal. We hope that adaptation of this method for *Neurospora* will be useful for many labs and the steps we changed from the *Ashbya* protocol can help researchers studying other filamentous fungi guide their attempts at adapting it for their systems.

## Acknowledgements

The authors wish to thank Samantha Dundon, PhD for contributions to early work surrounding adaptation of the smFISH protocol for *Neurospora.* We thank Andreia Verissimo, PhD of Dartmouth’s BioMT core for considerable assistance in establishing a protocol for chitinase expression and purification. Additionally, we acknowledge the efforts of Zuzana Burdikova, PhD for her work in assisting with early attempts at chitinase purification.

## Author Contributions

**Bradley M. Bartholomai:** Conceptualization, Methodology, Software, Validation, Formal analysis, Investigation, Data curation, Writing – Original Draft, Writing – Review & Editing, Visualization. **Amy S. Gladfelter:** Conceptualization, Methodology, Writing – Review & Editing, Supervision. **Jennifer J. Loros:** Conceptualization, Resources, Supervision, Project administration. **Jay C. Dunlap:** Conceptualization, Resources, Writing – Review & Editing, Supervision, Project administration, Funding acquisition.

---

**Supplementary Figure 1.**
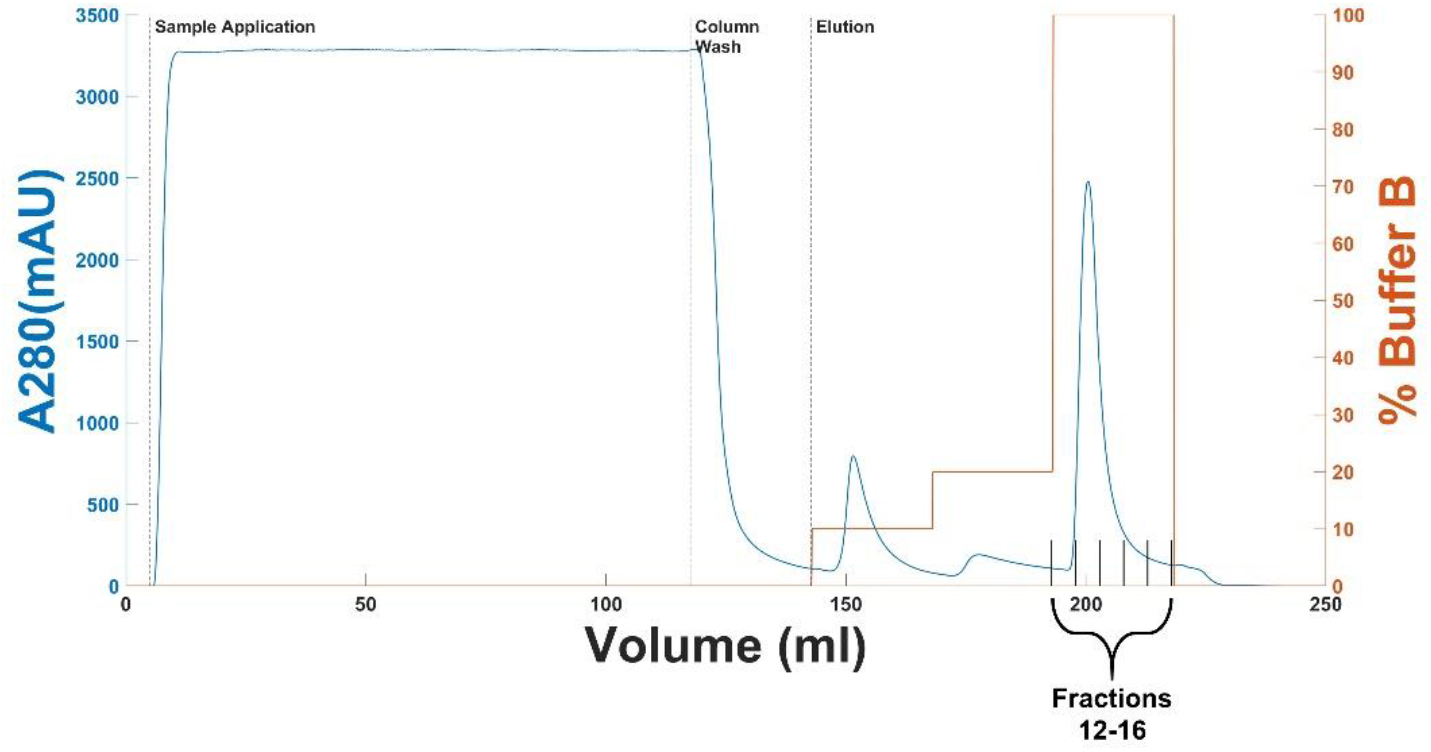
Chromatogram of NiNTA purification of Chitinase. Absorbance at 280 nm (blue) was measured throughout purification of chitinase of on a Cytiva Äkta FPLC. Sample application and pre-elution column washing was carried out with Buffer A, containing 20 mM imidazole. Elution was carried out stepwise with increasing concentration of Buffer B (orange), containing 500 mM imidazole. The eluate was collected in 1 column volume (5 ml) fractions. Fractions 12-16, corresponding to the greatest A280 peak and 100% buffer B were retained for further purification.

**Supplementary Figure 2.**
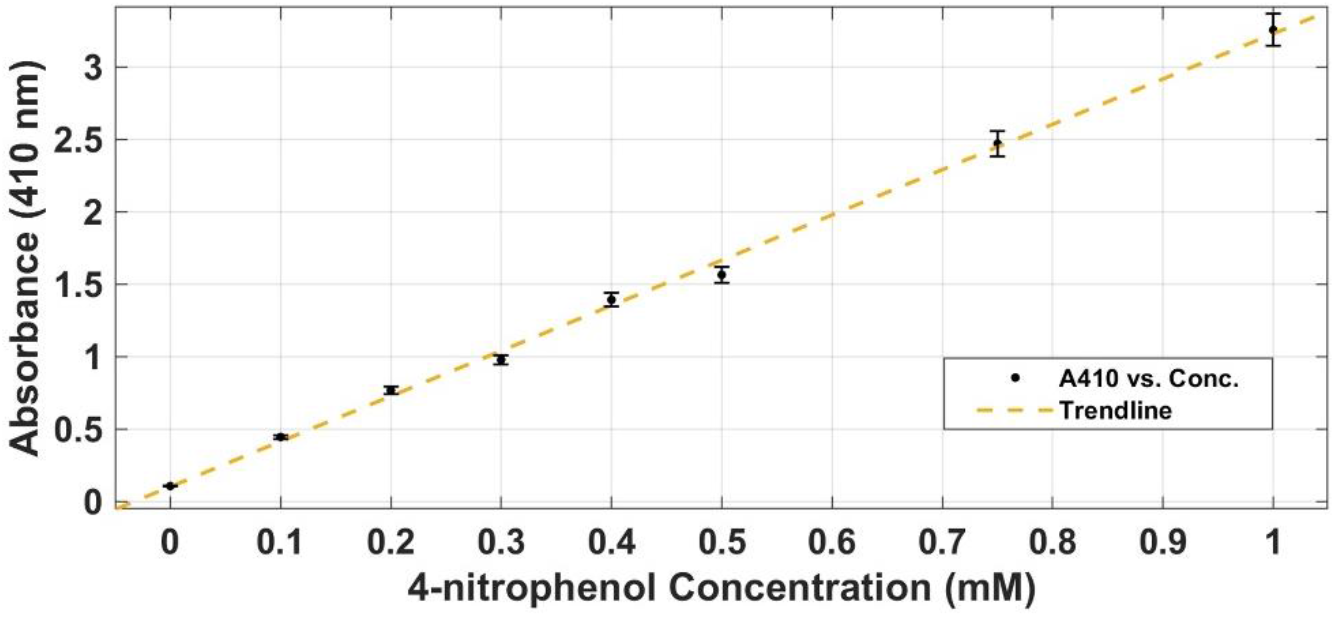
Standard curve of 4-nitrophenol. Linear regression was performed on a standard curve of known 4-nitrophenol concentrations measured on the same plate as experimental samples in a colorimetric chitinase assay using 4-Nitrophenyl β-D-N,N′,N″-triacetylchitotriose. These values were used to convert A410 values of 4-nitrophenol released in the chitinase assay to concentration values.

**Supplemental Figure 3.**
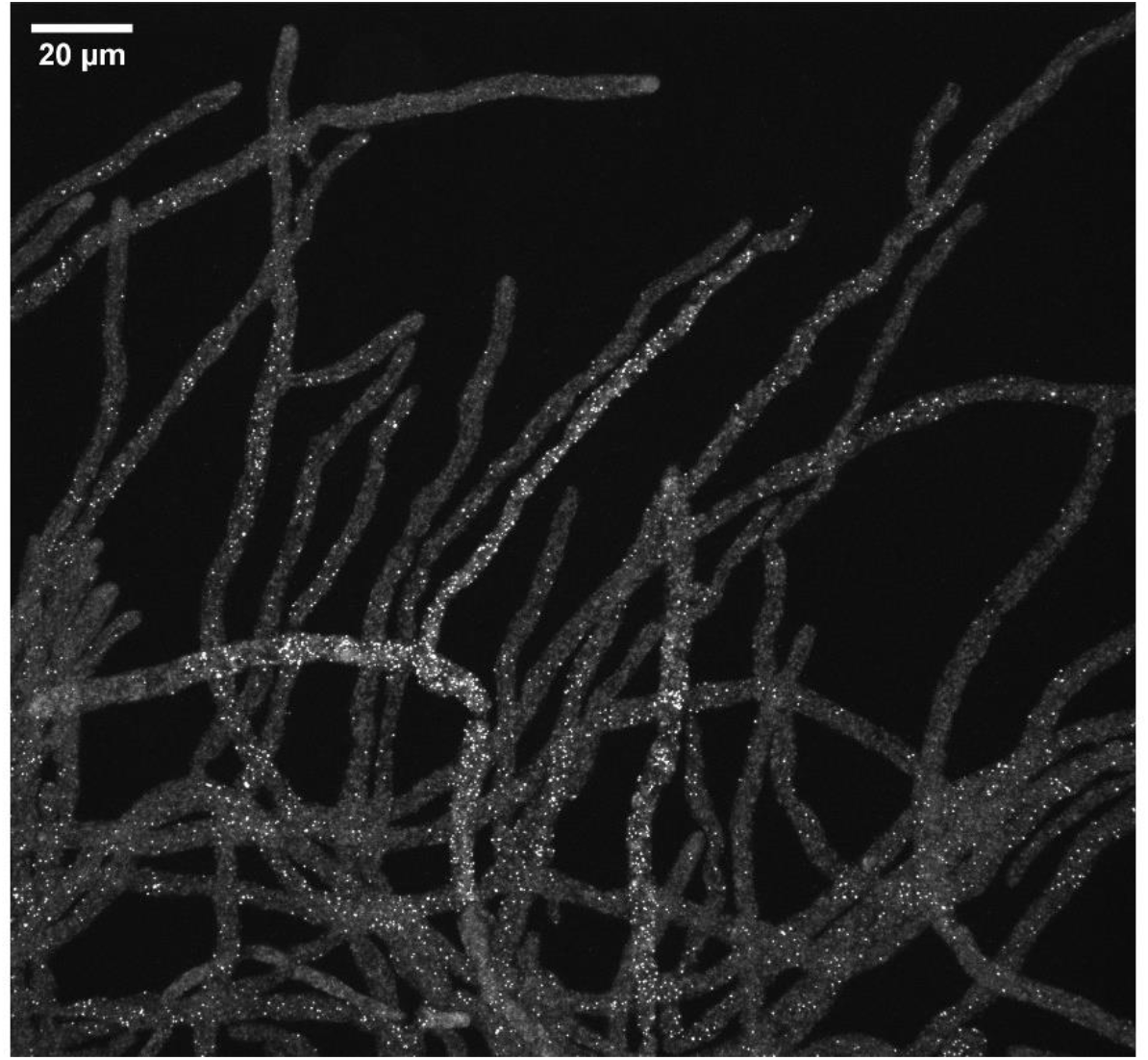
Heterogeneity between hyphae in a shared culture or mycelium is common in fungi. A maximum intensity z-projection of an uncropped field of view taken with a 2048 × 2048-pixel camera and 60X objective shows heterogeneous distribution of mRNA throughout a cluster of hyphae grown in a shared environment.

## References

Asmani, K.-L., et al., 2020. Biochemical and molecular characterization of an acido-thermostable endo-chitinase from Bacillus altitudinis KA15 for industrial degradation of chitinous waste. Carbohydrate Research. 495, 108089.

Baumann, S., et al., 2014. Endosomal transport of septin mRNA and protein indicates local translation on endosomes and is required for correct septin filamentation. EMBO reports. 15, 94–102.

Bolte, S., Cordelières, F. P., 2006. A guided tour into subcellular colocalization analysis in light microscopy. Journal of microscopy. 224, 213–232.

Bourett, T. M., et al., 1998. An improved method for affinity probe localization in whole cells of filamentous fungi. Fungal Genetics and Biology. 24, 3–13.

Caster, S. Z., et al., 2016. Circadian clock regulation of mRNA translation through eukaryotic elongation factor eEF-2. Proceedings of the National Academy of Sciences. 113, 9605–9610.

Craig, J. P., et al., 2015. Direct target network of the Neurospora crassa plant cell wall deconstruction regulators CLR-1, CLR-2, and XLR-1. MBio. 6, e01452–15.

Darken, M. A., 1961. Applications of fluorescent brighteners in biological techniques. Science. 133, 1704–1705.

de Bekker, C., et al., 2011. Single cell transcriptomics of neighboring hyphae of Aspergillus niger. Genome biology. 12, 1–12.

Dundon, S. E., et al., 2016. Clustered nuclei maintain autonomy and nucleocytoplasmic ratio control in a syncytium. Molecular biology of the cell. 27, 2000–2007.

Emerson, J. M., et al., 2015. period-1 encodes an ATP-dependent RNA helicase that influences nutritional compensation of the Neurospora circadian clock. Proceedings of the National Academy of Sciences. 112, 15707–15712.

Free, S. J., 2013. Fungal cell wall organization and biosynthesis. Advances in genetics. 81, 33–82.

Gasteiger, E., et al., 2005. Protein identification and analysis tools on the ExPASy server. The proteomics protocols handbook. 571–607.

Gomaa, E. Z., 2012. Chitinase production by Bacillus thuringiensis and Bacillus licheniformis: their potential in antifungal biocontrol. The Journal of Microbiology. 50, 103–111.

Harrington, B. J., Hageage Jr, G. J., 2003. Calcofluor white: a review of its uses and applications in clinical mycology and parasitology. Laboratory medicine. 34, 361–367.

Herold, I., Yarden, O., 2017. Regulation of Neurospora crassa cell wall remodeling via the cot-1 pathway is mediated by gul-1. Current genetics. 63, 145–159.

Hoch, H., et al., 2005. Two new fluorescent dyes applicable for visualization of fungal cell walls. Mycologia. 97, 580–588.

Hurley, J. M., et al., 2014. Analysis of clock-regulated genes in Neurospora reveals widespread posttranscriptional control of metabolic potential. Proceedings of the National Academy of Sciences. 111, 16995–17002.

Hurley, J. M., et al., 2018. Circadian proteomic analysis uncovers mechanisms of post-transcriptional regulation in metabolic pathways. Cell systems. 7, 613–626. e5.

Imdahl, F., Saliba, A.-E., 2020. Advances and challenges in single-cell RNA-seq of microbial communities. Current Opinion in Microbiology. 57, 102–110.

Kasuga, T., Glass, N. L., 2008. Dissecting colony development of Neurospora crassa using mRNA profiling and comparative genomics approaches. Eukaryotic Cell. 7, 1549–1564.

Kelliher, C. M., et al., 2020. Evaluating the circadian rhythm and response to glucose addition in dispersed growth cultures of Neurospora crassa. Fungal biology. 124, 398–406.

Laribi-Habchi, H., et al., 2015. Purification, characterization, and molecular cloning of an extracellular chitinase from Bacillus licheniformis stain LHH100 isolated from wastewater samples in Algeria. International Journal of Biological Macromolecules. 72, 1117–1128.

Lee, C., et al., 2016. Quantitative spatial analysis of transcripts in multinucleate cells using single-molecule FISH. Methods. 98, 124–133.

Lee, C., et al., 2013. Protein aggregation behavior regulates cyclin transcript localization and cell-cycle control. Developmental cell. 25, 572–584.

Li, G., Neuert, G., 2019. Multiplex RNA single molecule FISH of inducible mRNAs in single yeast cells. Scientific data. 6, 1–9.

Liu, H., et al., 2017. A-to-I RNA editing is developmentally regulated and generally adaptive for sexual reproduction in Neurospora crassa. Proceedings of the National Academy of Sciences. 114, E7756–E7765.

Lobo, M. D. P., et al., 2013. Expression and efficient secretion of a functional chitinase from Chromobacterium violaceum in Escherichia coli. BMC biotechnology. 13, 1–15.

Loros, J. J., 2020. Principles of the animal molecular clock learned from Neurospora. European Journal of Neuroscience. 51, 19–33.

Managadze, D., et al., 2010. Identification of PEX33, a novel component of the peroxisomal docking complex in the filamentous fungus Neurospora crassa. European journal of cell biology. 89, 955–964.

Mela, A. P., et al., 2020. Syncytia in Fungi. Cells. 9, 2255.

Menghiu, G., et al., 2019. Biochemical characterization of chitinase A from Bacillus licheniformis DSM8785 expressed in Pichia pastoris KM71H. Protein expression and purification. 154, 25–32.

Patel, P. K., Free, S. J., 2019. The genetics and biochemistry of cell wall structure and synthesis in Neurospora crassa, a model filamentous fungus. Frontiers in microbiology. 10, 2294.

Pieuchot, L., et al., 2015. Cellular subcompartments through cytoplasmic streaming. Developmental cell. 34, 410–420.

Raj, A., et al., 2008. Imaging individual mRNA molecules using multiple singly labeled probes. Nature methods. 5, 877–879.

Riquelme, M., et al., 2002. The effects of ropy-1 mutation on cytoplasmic organization and intracellular motility in mature hyphae of Neurospora crassa. Fungal Genetics and Biology. 37, 171–179.

Riquelme, M., et al., 2011. Architecture and development of the Neurospora crassa hypha–a model cell for polarized growth. Fungal biology. 115, 446–474.

Selker, E. U., Neurospora crassa. In: S. Maloy, K. Hughes, Eds.), Brenner’s Encyclopedia of Genetics (Second Edition). Academic Press, San Diego, 2013, pp. 61–63.

Songsiriritthigul, C., et al., 2010. Expression and characterization of Bacillus licheniformis chitinase (ChiA), suitable for bioconversion of chitin waste. Bioresource Technology. 101, 4096–4103.

Ursache, R., et al., 2018. A protocol for combining fluorescent proteins with histological stains for diverse cell wall components. The Plant Journal. 93, 399–412.

Verdín, J., et al., 2019. Off the wall: the rhyme and reason of Neurospora crassa hyphal morphogenesis. The Cell Surface. 5, 100020.

Vogel, H. J., 1956. A convenient growth medium for Neurospora (Medium N). Microb. Genet. Bull. 13, 42–43.

Wösten, H. A., et al., 2013. Heterogeneity in the mycelium: implications for the use of fungi as cell factories. Biotechnology letters. 35, 1155–1164.

Wu, C., et al., 2014. Genome-wide characterization of light-regulated genes in Neurospora crassa. G3: Genes, Genomes, Genetics. 4, 1731–1745.

Yamabhai, M., et al., 2008. Secretion of recombinant Bacillus hydrolytic enzymes using Escherichia coli expression systems. Journal of Biotechnology. 133, 50–57.

Zacchetti, B., et al., 2018. Multiscale heterogeneity in filamentous microbes. Biotechnology advances. 36, 2138–2149.

